# Inhibition of hyaluronan synthesis prevents β-cell loss in obesity-associated type 2 diabetes

**DOI:** 10.1101/2023.02.28.530522

**Authors:** Nadine Nagy, Gernot Kaber, Vivekananda G. Sunkari, Payton L. Marshall, Aviv Hargil, Hedwich F. Kuipers, Heather D. Ishak, Marika Bogdani, Rebecca L. Hull, Maria Grandoch, Jens W. Fischer, Tracey L. McLaughlin, Thomas N. Wight, Paul L. Bollyky

**Author notes:** Corresponding author: Paul. L. Bollyky, Stanford University School of Medicine, 279 Campus Drive, Beckman Center B241A, Stanford, CA, 94305. These authors contributed equally (Nadine Nagy, Gernot Kaber).

## Abstract

Pancreatic β-cell dysfunction and death are central to the pathogenesis of type 2 diabetes (T2D). We have identified a novel role for the inflammatory extracellular matrix polymer hyaluronan (HA) in this pathophysiology. Low levels of HA are present in healthy pancreatic islets. However, HA substantially accumulates in cadaveric islets of human T2D and islets of the db/db mouse model of T2D in response to hyperglycemia. Treatment with 4-methylumbelliferone (4-MU), an inhibitor of HA synthesis, or the deletion of the major HA receptor CD44, preserve glycemic control and insulin levels in db/db mice despite ongoing weight gain, indicating a critical role for this pathway in T2D pathogenesis. 4-MU treatment and the deletion of CD44 likewise preserve glycemic control in other settings of β-cell injury including streptozotocin treatment and islet transplantation. Mechanistically, we find that 4-MU increases the expression of the apoptosis inhibitor survivin, a downstream transcriptional target of CD44 dependent on HA/CD44 signaling, on β-cells such that caspase 3 activation does not result in β-cell apoptosis. These data indicate a role for HA accumulation in diabetes pathogenesis and suggest that it may be a viable target to ameliorate β-cell loss in T2D. These data are particularly exciting, because 4-MU is already an approved drug (also known as hymecromone), which could accelerate translation of these findings to clinical studies.

## INTRODUCTION

Type 2 diabetes (T2D) affects >34 million people in the U.S. alone^1^ and is associated with progressive β-cell dysfunction, insulin resistance, and ultimately β-cell demise^2, 3^. There is consequently great interest in identifying factors that impact β-cell health in T2D disease progression, so that these factors might be targeted therapeutically^4^.

The pathogenesis of β-cell dysfunction in T2D is complex and multifactorial. Prominent roles have been identified for host genetics^5^, amyloid deposition^6^, lipotoxicity^7^, and glucotoxicity^8^. The local inflammatory milieu is also thought to contribute to β-cell dysfunction^9^, leading to β-cell exhaustion and demise. However, the relevant tissue factors are unclear.

One factor that is abundant within inflamed tissues is the extracellular matrix (ECM) polymer hyaluronan (HA)^10^. HA accumulates within inflamed tissues in response to hyperglycemia^11, 12^, inflammatory cytokines^13^, and other triggers^10, 14, 15^, and promotes immune activation, cell migration and glycolytic control^16–18^. In T2D, HA is increased in serum, skeletal muscle and adipose tissue^19^. In healthy islets, modest amounts of HA are found within capillary walls and the peri-islet capsule, but intra-islet HA is low^17, 20^. HA was implicated in insulin resistance in peripheral tissues, particularly skeletal muscle^16, 21^.

Consistent with this role for HA in glycemic control, the HA receptor CD44 was implicated in the pathogenesis of T2D in a gene expression-based genome-wide association study (eGWAS) study incorporating 1,175 T2D case-control microarrays^22^. CD44^-/-^ mice were found to be resistant to diabetes induced using a high-fat diet while anti-CD44 antibody treatment of obese C57Bl6 (B6) mice decreased blood glucose levels and insulin tolerance^23, 24^. These effects were attributed to effects on insulin resistance^25, 26^ associated with CD44 expression on activated immune cells^27, 28^.

Together, these data implicate both HA and CD44 in insulin resistance associated with chronic inflammation. However, it is unclear from these studies whether this pathway is also relevant to the β-cell dysfunction and loss that is a prominent part of T2D pathogenesis. CD44 was linked to amino acid uptake for insulin synthesis^29^ and studies have examined HA-based scaffolds as a substrate for islet cell transplantation^30, 31^. However, the relevance of these studies to T2D have not previously been investigated.

Here, we have tested the hypothesis that HA and CD44 expression are increased in islets in T2D and that this pathway links the inflammatory milieu to β-cell dysfunction. We report that HA accumulates in both mouse and human islets in T2D and that this is associated with the progressive loss of β-cell mass and insulin production. Conversely, we find that inhibition of HA synthesis with 4-methylumbelliferone (4-MU) ^32–36^ is strongly protective against T2D. These results are particularly exciting given that 4-MU is already an approved drug called hymecromone. Our data suggest that it may be possible to repurpose 4-MU to treat and prevent β-cell loss in T2D.

## RESULTS

### HA is increased in islets of cadaveric organ donors with T2D

We first asked whether the finding of HA within inflamed tissues in response to hyperglycemia is observed in human pancreatic islets, and if this is relevant to human T2D. In order to do this, we obtained histologic sections of pancreatic tissues from human cadaveric donors with T2D and non-diabetic controls from the Juvenile Diabetes Research Foundation (JDRF) Network for Pancreatic Organ Donors with Diabetes (nPOD) program. Pancreatic sections from cadaveric donors with T2D (n=11) and non-diabetic controls (n=17) were stained for HA (Figure 1A, 1B). The characteristics of these donors are listed in Table 1. An average of 68 islets per subject with T2D and 119 islets per non-diabetic subject were analyzed in the tissue sections (Figure 1C). HA was observed predominantly around and along the peri- and intra-islet capillaries. This localization was similar in T2D and controls (Figure 1A, 1B).

**Figure 1.**
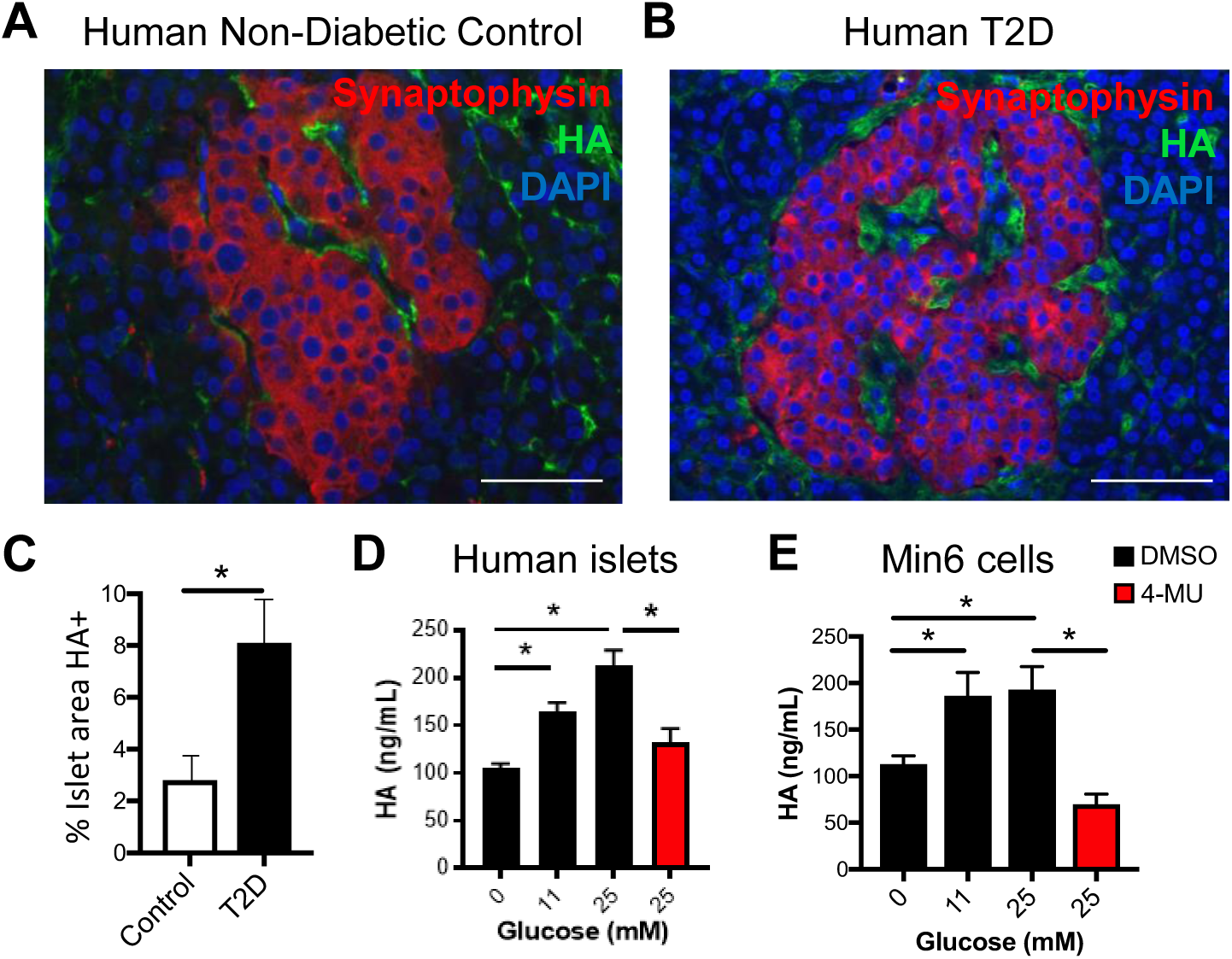
Islet HA is increased in human T2D. **A, B.** Representative staining of histologic islet sections from non-diabetic (control) (**A**) and Type 2 diabetic (T2D) (**B**) individuals stained for nuclei (DAPI, blue), HA (green), and synaptophysin (endocrine cell marker, red). Scale bars = 50 µm. A total of 750 T2D and 2030 control islets were analyzed. **C.** Percent positive islet area for HA. Data for **C** are mean ± SD of measurements performed in 17 non-diabetic and 11 T2D tissues. **D, E.** HA content in human islets and murine Min6 cells, a β-cell line, cultured in media containing glucose at the indicated concentrations, and with 50 µg/mL 4-MU added at the highest glucose concentration. Data in **D** and **E** are for triplicate measurements. * = p<0.05.

**Table 1.** Basic human patient characteristics.

However, the islet area positive for HA was significantly higher in T2D than in non-diabetic islets (Figure 1A-C). In contrast, the percentage of exocrine pancreas area positive for HA was not significantly different from controls (not shown), indicating that the increase in pancreatic HA content in these tissues is primarily in islets.

Next, we determined whether culture of human islets (Figure 1D) and Min6 cells (Figure 1E), a β-cell line, in media containing increased glucose concentrations resulted in increased HA concentration in the media. Indeed, the amount of HA released into the media increased proportionally to the concentration of glucose in the media (Figure 1D, 1E). Adding 4-MU, a pharmacologic inhibitor of HA synthesis^32–36^, to the media with the highest glucose concentration reduced the HA concentration significantly compared to the highest glucose concentration without 4-MU (Figure 1D, 1E). This was true for the isolated human islets as well as the Min6 cell line (Figure 1D, 1E).

Together these data indicate that HA is increased in human pancreas sections compared to non-diabetic controls, and that HA secretion is increased in human islets and β-cells in response to glucose.

### Intra-islet HA is increased in db/db mice and reduced upon oral 4-MU treatment

We next examined HA synthesis in the db/db mouse model of T2D. These C57Bl6 (B6) mice have a mutation in the long, signaling form, of the leptin receptor, resulting in progressive obesity and hyperglycemia.

To investigate the impact of HA inhibition on this model, we orally administered 4-MU to inhibit HA synthesis. Because HA is made by three different HA synthases (HAS1, HAS2, and HAS3) and the simultaneous constitutive deletion of all three is embryologically lethal, it was not practical to inhibit HA production using knock-out models.

We stained pancreatic islets of B6 and db/db mice for HA. We observed that HA was also present in islet capsule structures and intra-islet capillaries in both healthy and diabetic mice, as we reported previously^17, 19^. In diabetic db/db mice but not in normoglycemic conventional B6 controls, we found that HA deposits were present as punctate deposits (Figure 2A). The percentage of islet area that was positive for HA was significantly increased in db/db mice compared to B6 controls, and was significantly reduced in db/db mice which received 4-MU treatment (Figure 2B).

**Figure 2.**
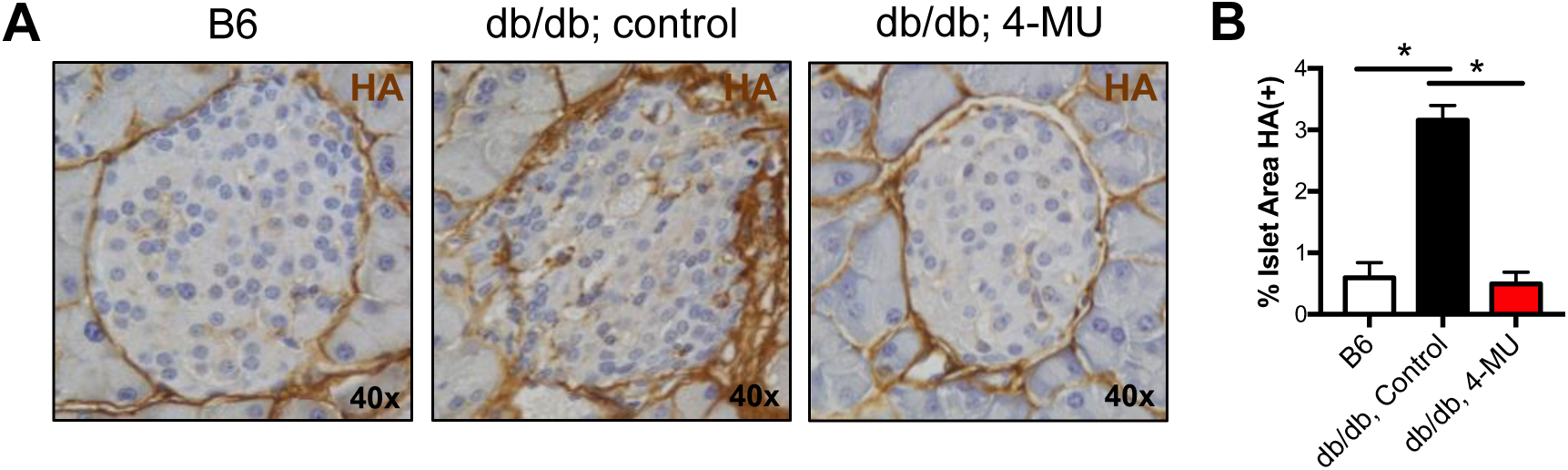
Diabetic db/db mice have increased islet HA. **A.** Representative HA staining of pancreatic islets from 15-week-old, male db/db mice that received 4-MU or control chow for 10 weeks starting at 5 weeks of age. Age- and gender matched non-diabetic B6 mice are shown for comparison. **B.** Quantified percent HA positive islet area in the same mice as in **A.**

### Inhibition of HA synthesis restores euglycemia in db/db mice

We next examined the impact of 4-MU on the blood glucose of conventional B6 and db/db mice. Conventional B6 mice started on 4-MU at 5 weeks of age had a non-significant BG reduction on 4-MU chow (Supplemental Figure 1A) and no improvement on intra peritoneal glucose tolerance test (IPGTT) responses at 12 weeks of age (Supplemental Figure 1B). The oral administration of 4-MU to male db/db mice starting at 5-6 weeks of age reliably decreased blood glucose (BG) levels compared to age and sex matched db/db mice fed control chow for 10 weeks (Figure 3A, 3B, 3D) despite the fact that these animals remained obese (Figure 3A, 3C, 3E). Heterozygous db/+ littermates, which don’t become diabetic and maintain normal weight, were included as internal control to the homozygous db/db mice (Figure 3A-E). The beneficial effect of 4-MU on glycemic control in the db/db mice was sustained over time. Consistent with this, fasting db/db mice fed 4-MU had a substantial improvement in BG upon IPGTT (Figure 3F). These data suggest that 4-MU reduces BG levels in diabetic mice with high HA level, but not in normoglycemic mice where HA level are lower at baseline.

**Figure 3.**
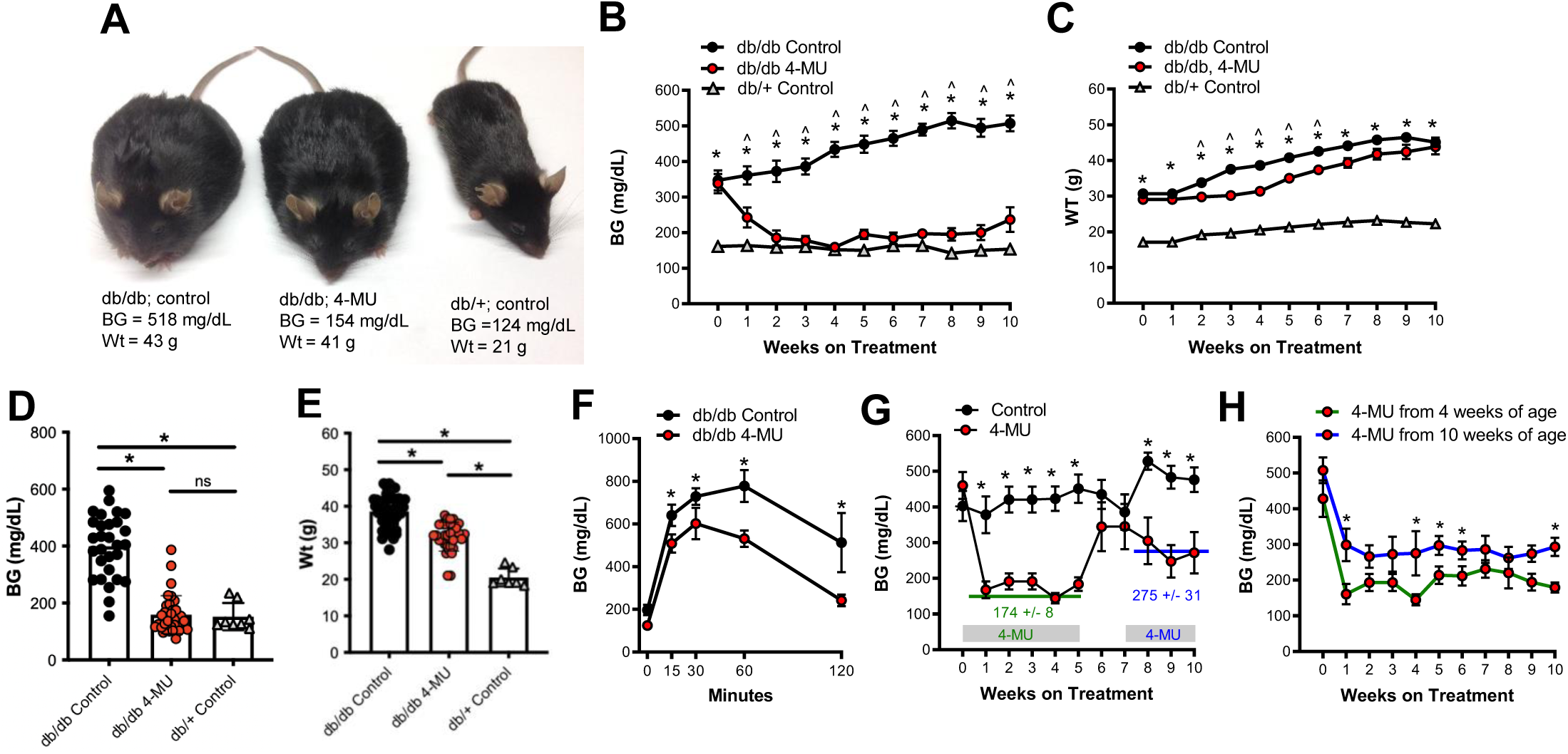
Oral 4-MU prevents hyperglycemia in obese db/db mice. **A-C.** Representative images (**A**), blood glucose (BG) values (**B**), and weights (**C**) over time for db/db mice treated with control chow, db/db mice treated with 4-MU chow as well as for db/+ heterozygous littermates treated with control chow. * = p<0.05 db/db control vs db/+ control. ^ = p<0.05 db/db control vs db/db 4-MU. n = >10 mice per group for each time point. **D, E.** Random (fed) BG values (**D**) and weights (**E**) for db/db mice treated with 4-MU or placebo for 4 weeks as well as their db/+ littermate controls on control chow. Each dot in **D** and **E** represents one mouse. Values over 600 mg/dL were excluded. **F.** IPGTT performed on fasting db/db mice treated with and without 4-MU chow. **G.** BG in mice fed 4-MU intermittently from 0-5 weeks and again from 7-10 weeks starting at 5 weeks of age. **H.** BG values over time for mice initiated on 4-MU chow at either 4 or 10 weeks of age. Individual time point comparisons were made using a student’s t-test. N = at least 8 mice per group for each figure panel. * = p<0.05.

One potential contributor to the observed glucose lowering could be an effect of 4-MU to reduce food intake and body weight. db/db mice on 4-MU chow indeed had an initial decrease in weight after the start of treatment but recovered within a few days (Figure 3C), as we have previously reported^35^. Consistent with this transient weight loss, db/db mice fed 4-MU decreased their food intake over the first week but by the second week had food intake comparable to chow-fed controls (Supplemental Figure 2). These data indicate that while decreased food intake could be a factor in 4-MU effects early upon initiation of this diet, this is unlikely to be an important factor over time.

A common approach to overcome the confounding effect of altered chow intake is drug delivery via oral gavage. However, because 4-MU is not soluble and has a short half-life in mice this approach was not feasible. Instead, we administered a soluble metabolite of 4-MU, 4-methylumbelliferyl glucuronide (4-MUG) to db/db mice in drinking water. We previously reported that 4-MUG is a bioactive metabolite of 4-MU that likewise inhibits HA synthesis^37^. As with 4-MU, db/db mice that consumed 4-MUG had a sustained decrease in BG (Supplemental Figure 3A, 3C), however, 4-MUG had no effect on body weight (Supplemental Figure 3B, 3D). Consistent with the glucose-lowering effects of 4-MUG, 4-MUG treatment also led to improved glucose tolerance upon IPGTT (Supplemental Figure 3E).

We conclude that while diminished caloric intake upon initial 4-MU treatment may prompt a transient drop in BG, the sustained improvement in glycemic control seen in db/db mice on either 4-MU or 4-MUG cannot be attributed to changes in body weight.

Together, these data indicate that inhibition of HA synthesis restores normoglycemia in diabetic db/db mice.

### 4-MU treatment is effective in mice with established diabetes

We next determined whether the timing of 4-MU treatment affected its efficacy to improve glycemic control of db/db mice. 4-MU treatment in 5-week-old db/db mice with established diabetes resulted in a significant improvement in BG over a 5-week period (Figure 3G). Withdrawal of 4-MU treatment resulted in an increase in BG to levels comparable to control db/db mice (Figure 3G). However, restarting 4- MU treatment in the same mice was again effective in decreasing BG, although this second treatment did not lower glucose to the same extent as the initial 4-MU treatment (Figure 3G).

In a separate experiment 4-MU treatment was initiated in diabetic db/db mice at 4- and at 10-weeks of age (Figure 3H). Both groups showed a significant decrease in BG within a week of 4-MU initiation. While lower BG were achieved when 4-MU treatment was initiated in 4-week-old mice, 4-MU was still highly effective at lowering BG when initiated at 10-weeks of age (Figure 3H).

Together these data indicate, that 4-MU treatment is effective in lowering BG in db/db mice with established diabetes and that while active treatment is required for glucose lowering, subsequent dosing of 4-MU remains effective at reducing BG.

### Islet CD44 expression is increased in db/db mice, and normalized upon HA synthesis inhibition

We next asked whether expression of CD44, the major HA receptor, was likewise increased in diabetic db/db mice. Using the same mice as in Figure 2A, we observed that CD44 immunoreactivity was increased in islets of db/db mice compared to B6 controls (Figure 4A), reflected by an increase in the percentage of islet area that was positive for CD44 (Figure 4B). 4-MU treatment resulted decreased islet CD44 (Figure 4A, 4B), suggesting that increased levels of islet CD44 may be driven by HA accumulation. Pericellular HA is known to stabilize CD44 on the cell surface in other systems^38^ and that may also be relevant here.

**Figure 4.**
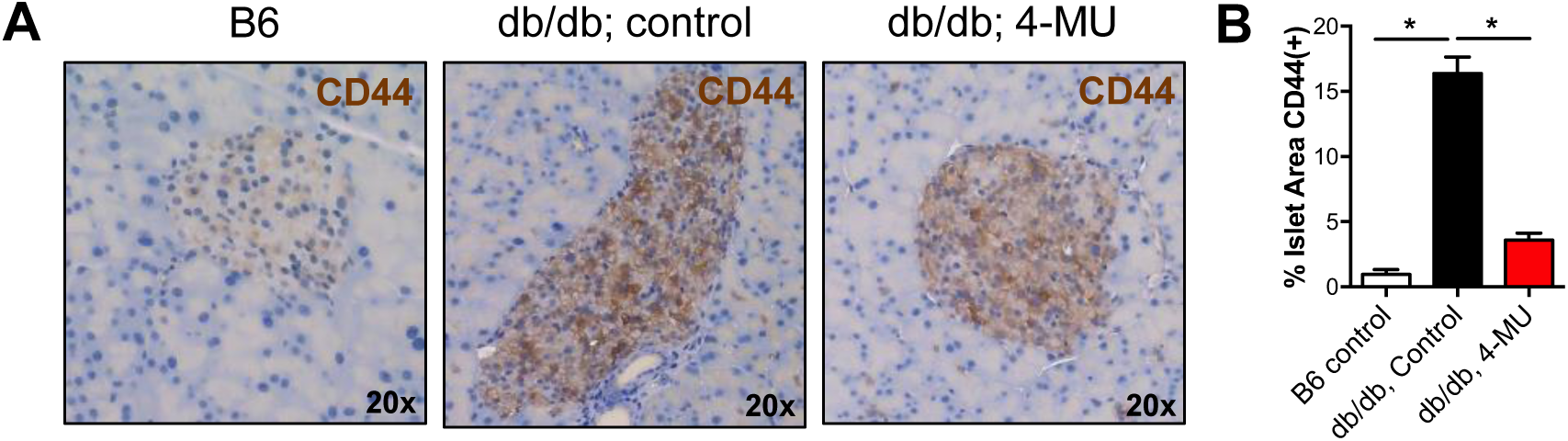
Obesity-associated diabetes is linked to increased islet CD44 in mice while 4-MU treatment reduces this. **A.** Representative CD44 staining of pancreatic islets from 15-week-old, male db/db mice that received 4-MU or control chow for 10 weeks starting from the age of 5 weeks. Age- and gender matched, non-diabetic B6 mice are shown for comparison. **B.** Quantified percent HA positive islet area in the same mice as in **A**.

### db/db.CD44^-/-^ mice do not develop diabetes

We next asked whether the major HA receptor CD44 is involved in glycemic control in db/db mice. To this end, we generated db/db mice lacking the CD44 receptor (db/db.CD44^-/-^ mice).

We observed that db/db.CD44^-/-^ mice are protected against developing hyperglycemia (Figure. 5A, 5B, 5D) despite being equivalently obese to db/db.CD44^+/+^ littermates (Figure 5A, 5C, 5E). Over time, db/db.CD44^-/-^ mice continued to gain weight but maintained comparable BG to their db/+.CD44^-/+^ non-diabetic littermates (Figure 5A-E). Upon IPGTT db/db.CD44^-/-^ mice had significantly better glucose tolerance compared to db/db.CD44^+/+^mice (Figure 5F). We also asked whether CD44 contributes to glycemic control in conventional B6 mice. B6.CD44^-/-^ mice had modestly decreased BG levels compared to strain, age, weight, and sex matched B6.CD44^+/+^ controls (Supplemental Figure 4A). We likewise observed modestly enhanced glycemic control in B6.CD44^-/-^ mice upon IPGTT (Supplemental Figure 4B). Together, these data indicate that CD44 is required for diabetes development in obese db/db mice and suggest that increased CD44 expression adversely impacts glycemic control.

**Figure 5.**
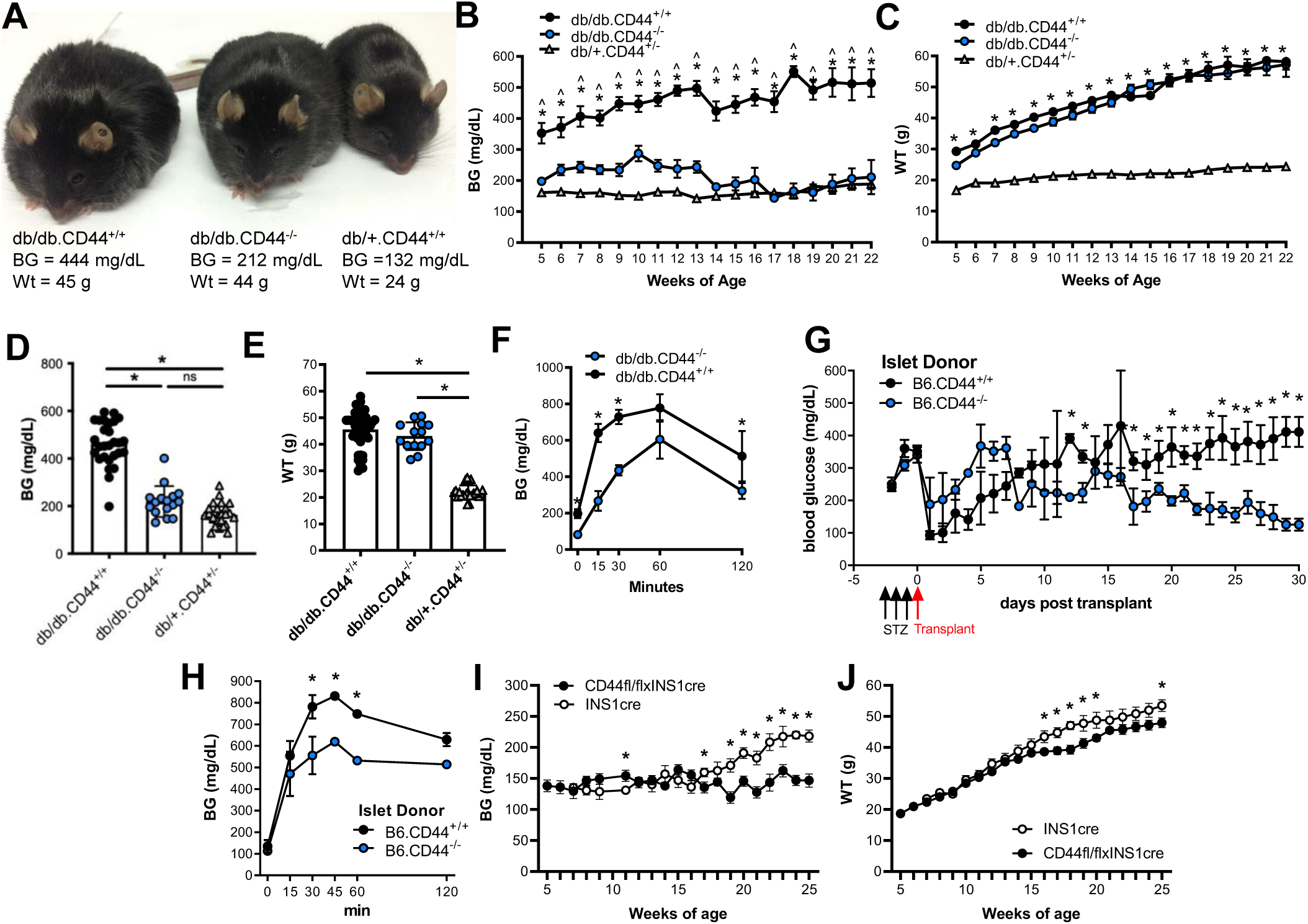
The absence of islet and β-cell CD44 protects obese db/db mice from diabetes. **A-C.** Representative images (**A**), blood glucose (BG) values (**B**), and weights (**C**) over time for db/db.CD44^+/+^ and db/db.CD44^-/-^ mice as well as for db/+ heterozygous littermates. * = p<0.05 db/db.CD44^+/+^ vs db/db.CD44^-/-^. ^ = p<0.05 db/db.CD44^+/+^ vs db/+.CD44^+/-^. **D, E.** Random (fed) BG values (**D**) and weights (**E**) for 12-week-old db/db.CD44^+/+^ and db/db.CD44^-/-^ as well as db/+ littermate controls. These are the same animals as in **A-C**. Each dot in **D**, **E**, represents one mouse. Values over 600 mg/dL were excluded. **F.** IPGTT performed on fasting db/db.CD44^+/+^ and db/db.CD44^-/-^ mice. **G.** BG levels following adoptive transfer of pancreatic islets harvested from B6.CD44^-/-^ or B6.CD44^+/+^ mouse and transferred into B6 mice previously rendered diabetic via STZ treatment. A timeline is shown on the X axis with black arrows marking STZ treatment and red arrows marking islet transplantation. **H.** IPGTT in the islet transplant mice from **G**. **I, J.** Random (fed) BG values (**I**) and weights (**J**) fromINS1cre (control) and CD44fl/flxINS1cre mice. n = at least 8 mice per group. * = p<0.05. Individual time point comparisons were made using a student’s t-test.

### Transplanted islets from B6.CD44^-/-^ mice correct diabetes longer in db/db/ mice than islets from B6.CD44^+/+^ mice

In order to determine whether the impact of CD44 on glycemic control was islet-specific in this model, we performed an islet transplantation experiment allowing us to specifically interrogate the role of islet CD44. Recipient B6.CD44^+/+^ mice were made diabetic via high dose streptozotocin (STZ), a β-cell-specific toxin, treatment over three days. On the fourth day, syngeneic islets from either B6.CD44^-/-^ or B6.CD44^+/+^ donor mice were transplanted under the kidney capsule into diabetic STZ-db/db.CD44^+/+^ recipients. We then followed these mice over the subsequent month (Figure 5G).

We observed an initial decline in glucose in all recipient mice, which may have been due to insulin release of some dying transplanted islets (Figure 5G). Initially glucose levels did not differ between groups, but starting around day 12 post-transplant, recipients of CD44^-/-^ islets exhibited improved and sustained glucose lowering (Figure 5G). Consistent with this, IPGTT testing revealed that STZ-db/db.CD44^+/+^ mice that received B6.CD44^-/-^ islets had better glycemic control in response to glucose challenge than mice that received B6.CD44^+/+^ islets (Figure 5H).

These data indicate that CD44 expression on islets contributes to the loss of glycemic control in this model, irrespective of CD44 expression in peripheral tissues.

### β-cell specific deletion of CD44 protects mice from becoming diabetic

To determine the direct impact of CD44 on β-cells and their glycemic control, we crossed CD44fl/fl with INS1cre mice, to generate β-cell specific CD44 deletion, and fed these mice a high fat diet. BG of the control mice on the high fat diet increased over time, whereas the BG of the CD44fl/flxINS1cre mice stayed stable (Figure 5I). CD44fl/flxINS1cre mice showed a modest increase in body weight relative to INS1cre controls after 16 weeks of age (Figure 5J).

These data support our previous finding, that limiting β-cell CD44 expression is an effective strategy for maintaining glycemic control in mice.

### 4-MU partially restores islet insulin positive area and reduces islet apoptosis in db/db mice

Given these effects on glycemic control, we next asked whether 4-MU and CD44 impact insulin staining on histology. We observed that 16-week-old db/db mice on long term (10 weeks) 4-MU treatment had a substantially greater islet insulin positive area than mice that received control chow, though this was nonetheless still reduced compared to sex-, and age-matched B6 controls (Figure 6A, 6B). Islet insulin area was likewise increased in db/db.CD44^-/-^ relative to db/db.CD44^+/+^ mice (Figure 6C, 6D), in this case insulin positive area improved to a level comparable to db/+ control mice.

**Figure 6.**
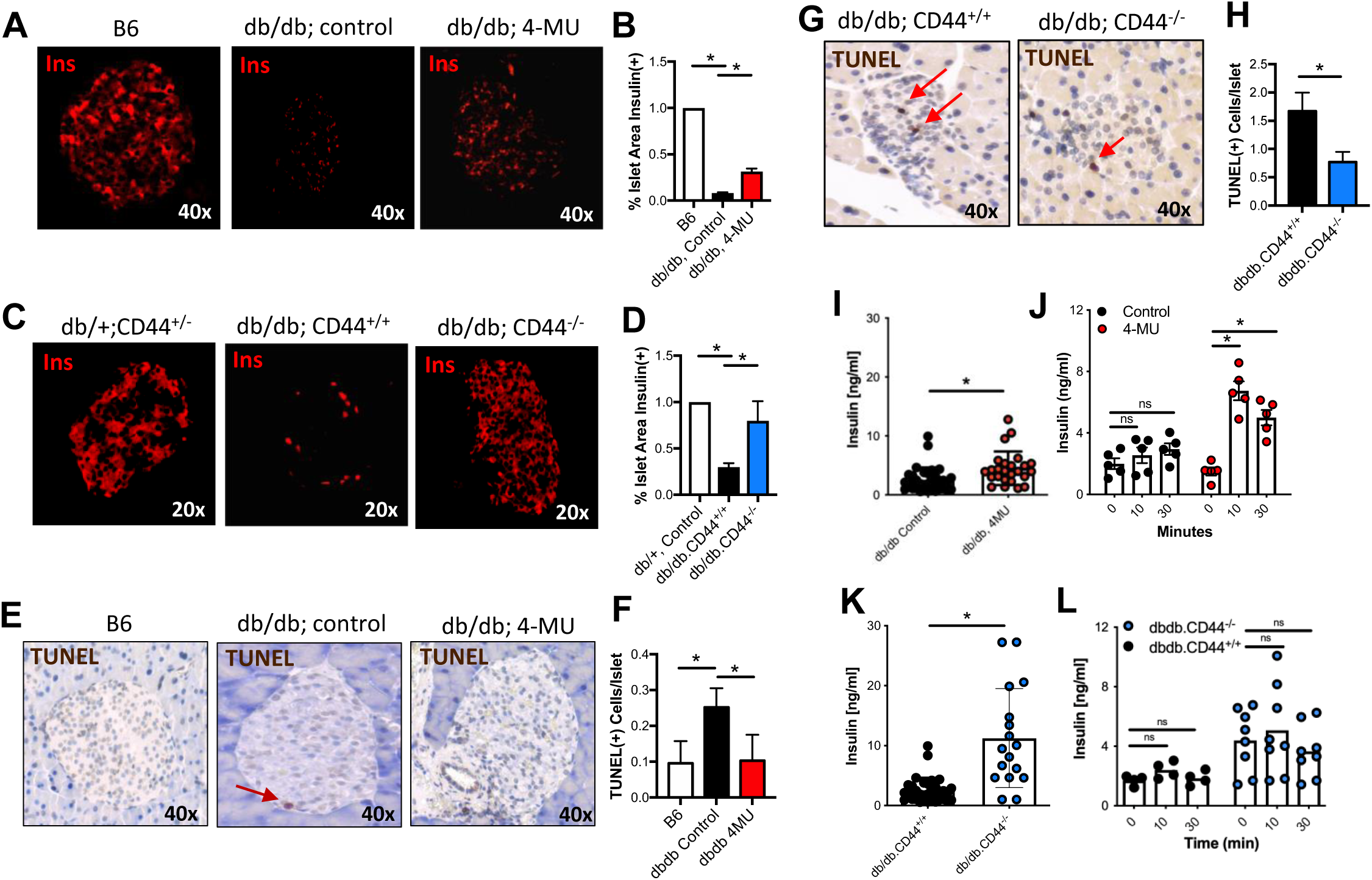
Inhibition of HA synthesis preserves islet insulin staining and prevents islet cell death in db/db mice. **A.** Representative insulin staining of pancreatic islets from db/db mice, treated with control of 4-MU diet**. B.** Quantified percent insulin positive islet area in the same mice as in **A. C.** Representative insulin staining of pancreatic islets from of pancreatic islets from db/db.CD44^+/+^ and db/db.CD44^-/-^ mice. **D.** Quantified percent insulin positive islet area in the same mice as in **C. E.** Representative TUNEL staining of pancreatic islets from the same mice as in **A**. **F.** Quantified percent TUNEL positive islet area in the same mice as in **E. G**. Representative TUNEL staining of pancreatic islets from db/db.CD44^+/+^ and db/db.CD44^-/-^ mice. **H**. Quantified percent TUNEL positive islet area in the same mice as in **G.** For all studies, at least 25 islets per mouse and at least 4 animals were examined per condition. **I, J.** Random (fed) (**I**) and fasting (**J**) serum insulin levels in db/db mice treated with and without 4-MU chow. **K, L.** Random (fed) (**K**) and fasting (**L**) serum insulin levels in db/db.CD44^+/+^ and db/db.CD44^-/-^ mice. **I-L.** Data are representative of 3 independent experiments. * = p<0.05.

In light of these data, we wondered whether 4-MU might influence β-cell apoptosis. Indeed, db/db mice on 4-MU also exhibited a reduction in TUNEL positive cells upon histologic staining (Figure 6E, 6F). The same could be observed in db/db.CD44^-/-^ mice compared to regular db/db mice (Figure 6G, 6H). Of note, the relatively low number of apoptotic cells relative to the total decline in β-cell mass in these animals probably reflects the fact that apoptotic β-cells are rapidly cleared by phagocytosis^27^. Therefore, only cells that are apoptotic immediately at the fixation time are counted (counting of apoptotic β-cells in many islets compensate for this low number).

Together, these data indicate that HA reduction through 4-MU treatment or deleted CD44 expression halts the progressive decline in islet insulin positive area typically seen in db/db mice in association with reduced apoptosis.

### 4-MU treatment or deletion of CD44 promotes insulin secretion in db/db mice

To determine whether the improved glucose tolerance is explained, at least in part, by improved insulin release, we treated mice with 4-MU and measured their insulin secretion in response to glucose stimulation. Non-fasted serum insulin levels were indeed increased in db/db mice treated with 4- MU (Figure 6I). Moreover, insulin release was significantly increased at 10 and 30 min after i.p. glucose in db/db mice maintained on 4-MU, in contrast to db/db mice fed conventional chow which had a minimal insulin response to i.p. glucose (Figure 6J), consistent with severe β-cell dysfunction in this model. These data indicate that 4-MU treatment promotes insulin secretion in db/db mice. These data indicate that 4-MU treatment promotes insulin secretion in db/db mice. Loss of CD44 in db/db mice was also associated with increased insulin levels, although db/db.CD44^-/-^ mice did not exhibit a greater insulin response to glucose compared to db/db.CD44^+/+^ mice (Figure 6K, 6L).

Together, these data indicate that 4-MU and the absence of CD44 improve basal (fed) insulin levels in vivo and that 4-MU treatment preserves the ability to secrete insulin in response to glucose challenge.

Consistent with these findings, we observed that insulin mRNA expression was decreased in Min6 cells engineered to overexpress CD44 (Min6.CD44) compared to regular Min6 cells (Min6.Luc) (Supplemental Figure 5A). HA treatment likewise suppressed insulin mRNA production by Min6 cells (Supplemental Figure 5B). Conversely, 4-MU treatment did not increase insulin mRNA and did not overcome the effects of exogenous HA (Supplemental Figure 5B). These data indicate that HA and CD44 influence the expression of insulin mRNA.

Overall, these functional and histologic data are perhaps most consistent with effects of HA/CD44 on multiple aspects of β-cell stability, insulin expression and insulin secretion. Since β- cell loss obviously impacts β-cell function, it is unfortunately not possible to unravel the precise extent of these different contributions.

### 4-MU protects β-cells from STZ induced cell death, but does not decrease caspase levels

We asked whether the impact of these pathways on β-cell function and stability were relevant to other forms of injury. To this end, we administered STZ, to healthy B6 mice. In particular, we administered what we have termed “very low dose STZ” − 40 mg/kg for 4 consecutive days. We previously reported that this regimen generates insulitis and transient (< 4 weeks) hyperglycemia in B6 mice^39^.

We find that very-low dose STZ treatment regimen results in intra-islet deposition of HA (Figure 7A) and an increase in total islet HA content (Figure 7B). We treated B6 mice with 4-MU and STZ and find that 4-MU treatment protected mice from the effects of very low dose STZ; mice maintained on the drug for two weeks experienced less of an increase in BG upon low dose STZ treatment (Figure 7C). We also observed, that B6 mice with or without 4-MU treatment develop diabetes to an equivalent extent upon treatment with high dose STZ (200 mg/kg for 3 consecutive days), a regimen that causes β-cell destruction and terminal diabetes^40^ (Figure 7D).

**Figure 7.**
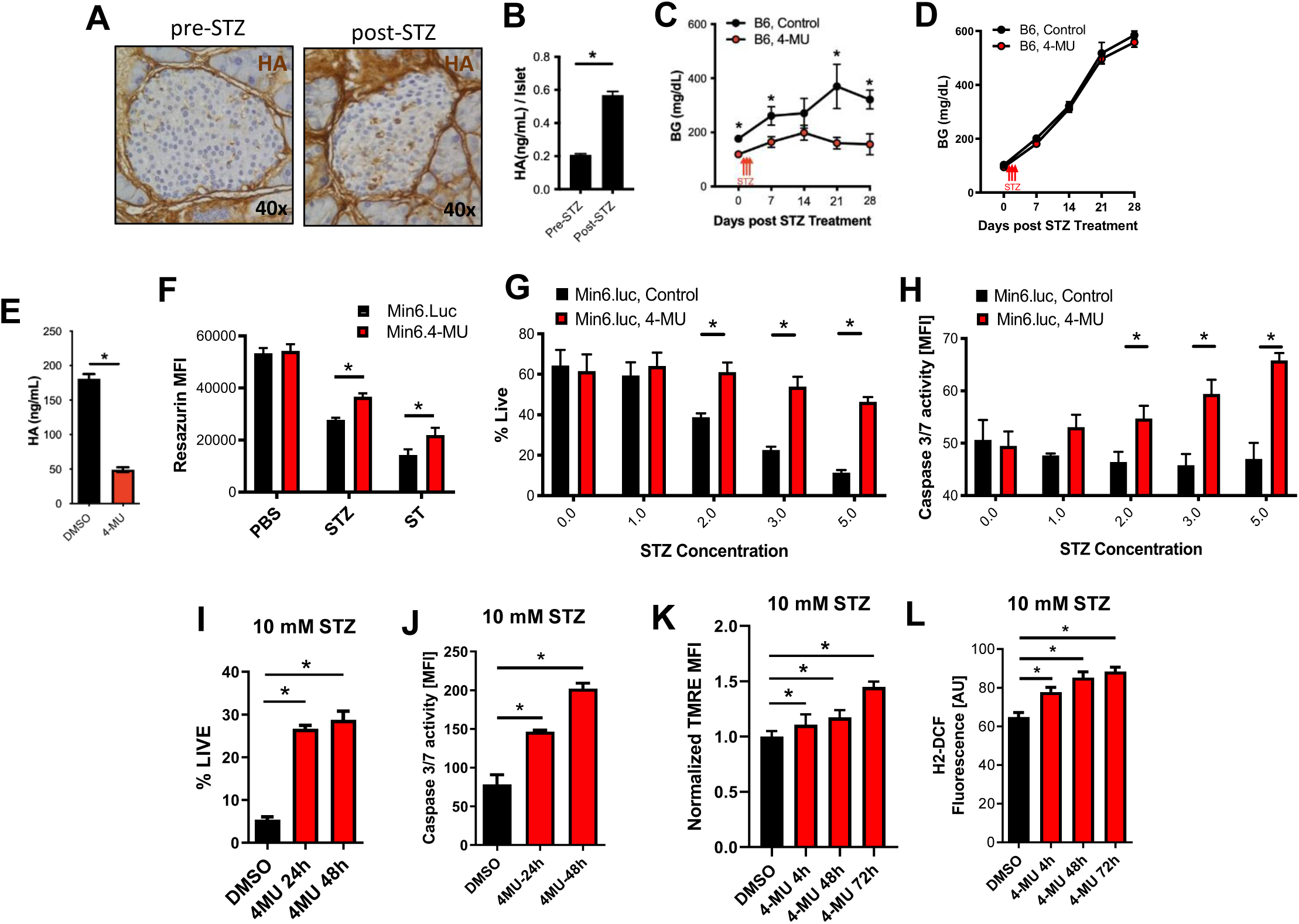
4-MU protects β-cells from cell death through low-dose STZ but does not decrease caspase levels. **A.** Representative im*a*ges of islet cells pre and post STZ treatment stained for HA (brown). **B.** Quantification of HA in ng/mL per islet from pre and post treatment with STZ, a β-cell-specific toxin. We administered low dose STZ (40 mg/kg) for 4 consecutive days – a regimen previously shown to cause insulitis and transient hyperglycemia in B6 mice. **C.** Random (fed) BG measurements post low dose STZ (40 mg/kg) treatment of B6 mice either control or 4-MU chow. **D.** Random (fed) BG measurements post high dose STZ (200 mg/kg) treatment of B6 mice either control or 4-MU chow. N = 6-10 mice per group. **E**. HA content of Min6 cells treated with 4-MU or DMSO as control. **F**. Min6. cells were treated with streptozotozin (STZ), staurosporin (ST) or PBS as control and resazurin MFI as a readout for live cells was measured. 4-MU was added to those cell lines and treatments as described in **F** and resazurin MFI was measured. **G**. % live cells measured in Min6 cells after STZ treatment at the indicated concentrations. **H**. Caspase 3/7 activity measured in the same cells and treatments as in **F**. **I-L.** Min6 cells pre-treated with 4-MU for 24 hrs before 10 mM STZ was administered. In this experimental setting % live cells (**I**), caspase 3/7 activity (**J**), TMRE (**K**) and H2-DCF (**L**) was measured. **G**, **H** and **I**, **J** are paired data from single experiments. * = p<0.05

### 4-MU prevents β-cell apoptosis

We next asked whether this pathway impacts β-cell apoptosis. We found that HA production by Min6 could be suppressed using 4-MU (Figure 7E), and that at baseline cell viability of identical numbers of Min6 cells with and without 4-MU treatment was equivalent (Figure 7F). 4-MU treatment improved cell viability upon treatment with either streptozotocin (STZ) or another pro-apoptotic agent staurosporine (ST) (Figure 7F). This was the case across a range of STZ concentrations, measured as percent live cells (Figure 7G). Of note, neither CD44 expression nor 4-MU treatment altered expression of GLUT-2, the receptor for STZ (Supplemental Figure 6A, 6B) indicating that receptor availability was not responsible for the effects on STZ toxicity seen here.

### 4-MU protects β-cells from STZ-induced cell death

Despite their enhanced viability (Figure 7G), Min6.luc cells treated with 4-MU nonetheless exhibited increased caspase3/7 activity compared to Min6.luc control treated cells over a range of STZ concentrations (Figure 7H). This was likewise the case over a range of time points of 4-MU treatment (Figure 7I and 7J). These data indicate that despite remaining viable, 4-MU treated Min6 cells nonetheless demonstrate initiation of what is typically a pro-apoptotic mechanism.

Consistent with this, we found that Min6 cells exhibited depressed mitochondrial membrane potential, as measured by tetramethylrhodamine, ethyl ester (TMRE) levels while 4- MU treatment restores this (Figure 7K). 4-MU treatment likewise increased expression of H2- DCF, an indicator of mitochondrial oxidative stress (Figure 7L).

These data suggest that STZ treatment triggers pro-apoptotic pathways in these cells but that 4-MU treatment prevents apoptosis and preserves β-cell mitochondria number and function.

### Absence of CD44 is protective against very low-dose STZ treatment

Consistent with these results with 4-MU treatment, we likewise observed that B6.CD44^-/-^ mice developed less severe hyperglycemia than B6.CD44^+/+^ mice following very low dose STZ treatment (Figure 8A). Glucose tolerance was also improved, as shown in an IPGTT performed on day 21 after low dose (40 mg/kg) STZ treatment (Supplemental Figure 7A). B6.CD44^-/-^ mice develop diabetes to an equivalent extent to B6.CD44^+/+^ mice upon treatment with a high dose (200 mg/kg) STZ protocol (Supplemental Figure 7B) indicating that these animals are only relatively and not absolutely resistant to STZ.

**Figure 8.**
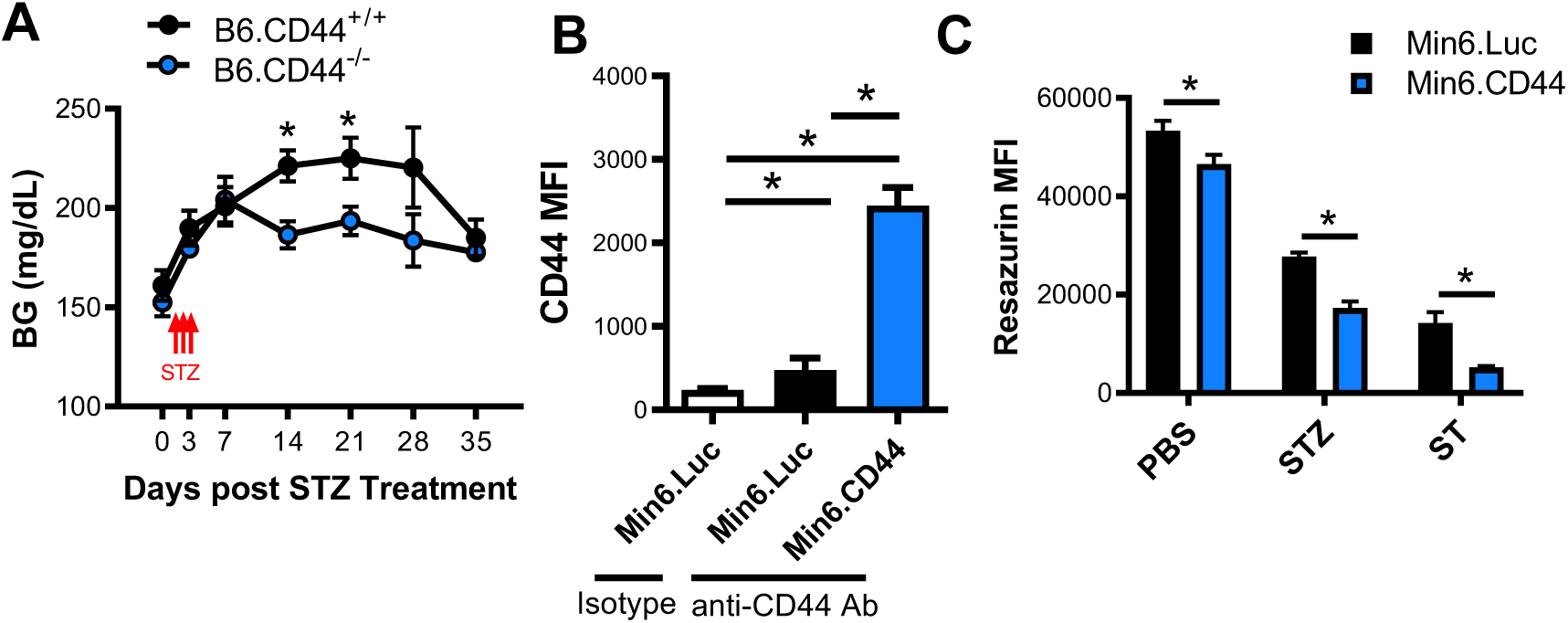
Absence of CD44 is protective against low-dose STZ treatment. **A.** We administered low dose STZ (40 mg/kg) for 4 consecutive days – a regimen previously shown to cause insulitis and transient hyperglycemia in B6 mice to db/db.CD44^+/+^ and db/db.CD44^-/-^ mice and measured random (fed) BG for up to 35 days post low dose STZ treatment. **B**. CD44 MFI levels on Min6.luc and Min6.CD44 cells. **C**. These same cells treated with streptozotozin (STZ), staurosporin (ST) or PBS as control and resazurin MFI as a readout for live cells was measured.

To better understand how this pathway impacts β-cells we studied Min6 cells, a β-cell line that normally expresses minimal CD44 at baseline. These cells were engineered to express high levels of CD44 (Min6.CD44) or a luciferase control (Min6.Luc) (Figure 8B).

We next asked whether this pathway impacts β-cell apoptosis. We found that at baseline identical numbers of Min6.CD44 cells were only 86% as viable as Min6.Luc cells (Figure 8C). Upon challenge with agents that induce apoptosis, Min6.CD44 cells were only 62% as viable as Min6.Luc cells upon streptozotocin (STZ) treatment and only 37% as viable upon staurosporine (ST) treatment (Figure 8C).

These data suggest that CD44 adversely impacts β-cell viability when exposed to STZ.

### Survivin is linked to HA and β-cell health

Given our finding that STZ treatment triggers caspase 3/7 but not apoptosis, we considered a potential role for survivin. Survivin, is a downstream transcriptional target of CD44, dependent on HA/CD44 signaling, and a regulator of caspase 3/7 activity that can prevent cell death. Survivin levels have been implicated in β-cell viability.

We observed that islets from db/db mice on long term (10 weeks) 4-MU treatment had a substantially greater islet survivin positive area than mice that received control chow (Figure 9A, 9B). The islet survivin positive area was likewise increased in db/db.CD44^-/-^ relative to db/db mice (Figure 9A, 9B).

**Figure 9.**
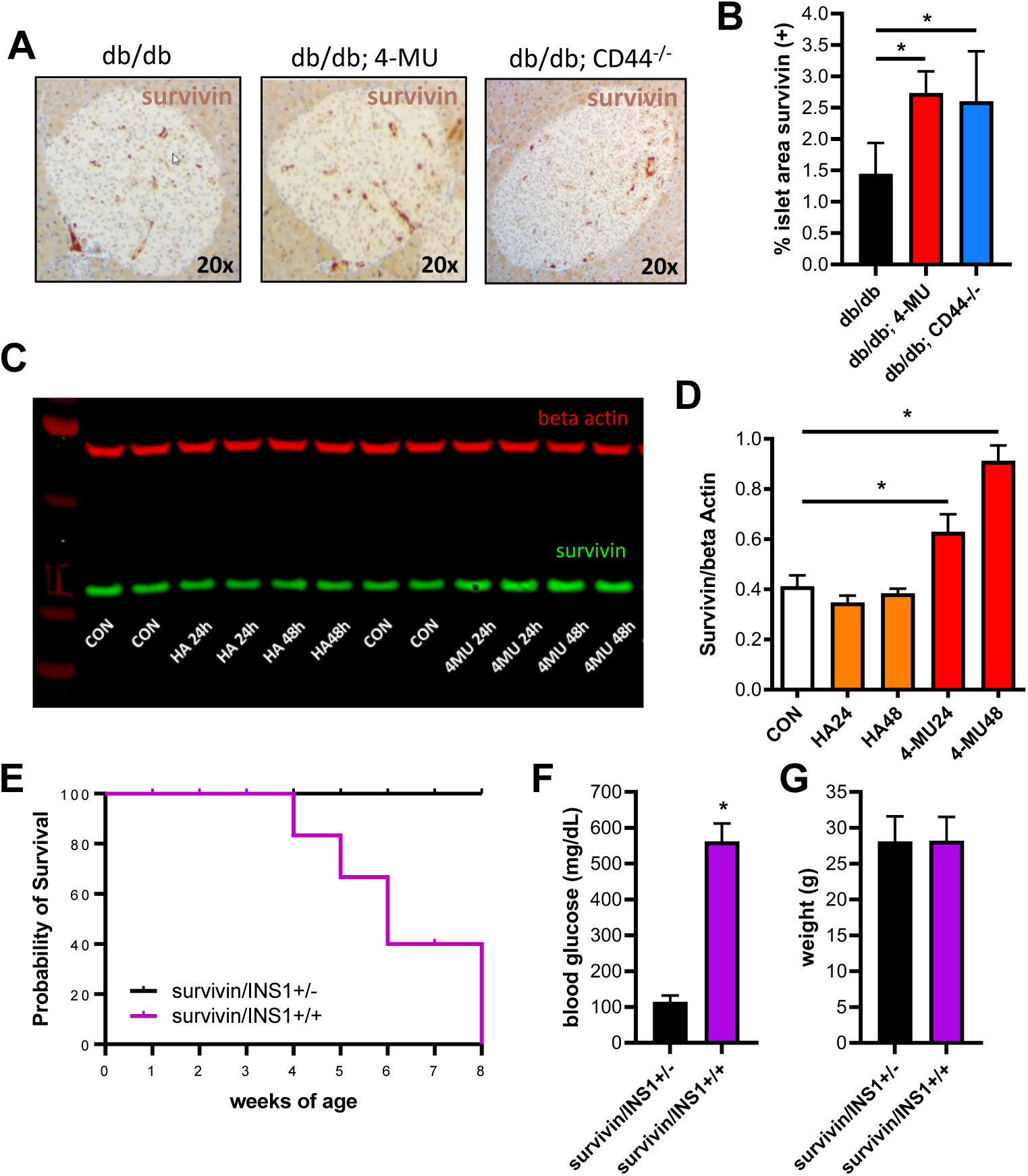
Survivin is linked to HA and beta cell health. **A.** Representative survivin staining of pancreatic islets from 15-20-week-old, male db/db mice that received 4-MU or db/db.CD44^-/-^ mice. **B.** Quantified percent survivin positive islet area in the same mice as in **A. C.** Survivin expression of Min6 cells treated with HA and 4-MU for 24 and 48 hours. **D**. Quantified survivin expression of Min6 in WB in **C**. **E**. Kaplan-Meyer survival curve for survivinfl/fl/INS1cre homozygous mice and survivinfl/fl/INS1cre heterozygous mice. **F, G**. Random (fed) BG values (**F**) and weights (**G**) of survivinfl/fl/INS1cre homozygous mice and survivinfl/fl/INS1cre heterozygous mice. n = 8-12 mice per group. * = p<0.05.

To better understand how this pathway impacts β-cells we studied Min6 cells, a β-cell line that normally expresses minimal CD44 at baseline. Via Western blot we detected survivin protein in Min6 cells treated with HA or 4-MU for 24 hrs or 48 hrs, using beta actin as loading control (Figure 9C). Survivin expression was slightly reduced in cells treated with HA, while 4-MU treatment significantly increased survivin expression in the Min6 cells (Figure 9D). These data indicate that 4-MU treatment induces survivin expression in vivo and in vitro.

To investigate the in vivo impact of β-cell survivin, we generated transgenic survivinfl/flxINS1cre (survivin/INS1) mice, in which survivin is knocked-out on exclusively on β- cells. The homozygous survivin/INS1 mice showed poor survival, most mice died within a few weeks after weaning (Figure 9E). We observed that those mice had severe hyperglycemia (Figure 9F), while their body weight (Figure 9G) was comparable to their heterozygous littermates. These data indicate that survivin plays a critical role in β-cell survival.

## DISCUSSION

We report that HA accumulation contributes to hyperglycemia via β-cell dysfunction and loss that characterizes T2D. We find that HA is increased within the pancreatic islets of both T2D patients and the db/db model of the disease. Conversely, inhibition of HA synthesis or absence of the HA receptor CD44 preserves glycemic control without weight loss. Together, our data indicate that the HA/CD44 pathway is a critical link between obesity and diabetes.

Our data implicate HA and CD44 in the loss of β-cells associated with T2D. Both inhibition of HA synthesis with 4-MU/4-MUG or the absence of CD44 preserved insulin staining and secretion in db/db mice despite long-term obesity. We propose a model whereby islet HA is produced by multiple cell types within islets in response to pro-inflammatory cues, including hyperglycemia^11^, inflammatory cytokines^13^, toxins, or other stressors^26, 27^. CD44 signaling likewise increased in T2D and, if HA is also present, increases β-cell apoptosis, leading to the progressive loss of β-cell mass seen in T2D. Conversely, targeting either HA or CD44 may interrupt this pathway and preserve glycemic control.

HA and CD44 influence the susceptibility of β-cells to apoptotic death. 4-MU and the absence of CD44 protected mice from very low dose STZ (but not from high dose STZ). Similarly, 4-MU and the absence of CD44 also protected β-cell from apoptosis in vitro, despite the upregulation of caspase 3/7. These data are consistent with established roles for HA and CD44 in cellular homeostasis pathways in other systems. CD44 is linked to both apoptosis and survival via multiple cell-type dependent mechanisms^41–45^.

We find that 4-MU treatment is associated with increased levels of survivin, a molecule that regulates the pro-apoptotic activity of caspases, thereby leading to negative regulation of apoptosis or programmed cell death^46^. Evidence suggests that survivin can inhibit both the extrinsic and intrinsic (mitochondrial) pathways of programmed cell death by blocking the activity of several caspase proteins^47^. Studies about survivin in mice have shown that perinatal survivin is essential for pancreatic β-cell homeostasis and that the postnatal expansion of the pancreatic β-cell mass is dependent on survivin^48^. Another study could show that the inhibition of apoptosis by survivin improves islet transplantation outcomes in mice^49^. Survivin is known to be a downstream transcriptional target of CD44^50^, dependent on HA/CD44 signaling, which is playing an important role in apoptosis^49^. Furthermore, a novel study using microarray analysis highlighted the relationship of HA/CD44 in activating the transcription of survivin as underlying signaling mechanism^50^. These data fit very well with our data suggesting that survivin plays an important role in β-cell survival and that HA is an important regulator of survivin.

When one considers that mice treated with 4-MU are also resistant to autoimmune type 1 diabetes (T1D)^36^ and that islet HA deposits and CD44 also accumulate in that disease^20^, inhibiting HA synthesis may protect islets in fundamental ways that transcend the differences between hyperglycemic glucotoxicity, oxidative stress due to STZ treatment, and cytotoxic killing.

These data do not preclude roles for HA and CD44 in insulin resistance. Previous reports have clearly shown that this pathway affects insulin sensitivity in obese animals and humans, as discussed in the introduction and reviewed elsewhere^11, 12^. Nonetheless, many of our assays and histologic studies demonstrate additional islet and β-cell-specific effects of HA and CD44. Together, these effects on both islet pathophysiology and insulin sensitivity point to important, overlapping roles for HA/CD44 responses in diabetes.

These data raise several questions for future studies. The cellular source of HA production in T2D islets remains unknown. Hyperglycemia has been reported to induce shedding of the HA rich glycocalyx of endothelium^51^ and multiple inflammatory cell types are known to produce HA in response to glucose. Which cell(s) are the source of HA in the islets awaits further investigation. Finally, numerous other extracellular matrix components (e.g. hyaladherins) are known to be present in islets and to interact with HA and its binding partners^36, 52–54^. The contributions of these molecules to CD44 signaling and islet physiology remain incompletely understood and await further investigation.

It may be possible to target this HA synthesis to preserve β-cell mass in T2D. 4-MU is already an established therapy approved in humans. Sold as “Hymecromone” under a variety of trade names, 4-MU is used throughout Europe and Asia (but not the USA) to prevent gallstones. It has been used for >30 years in both children and adults, has well established pharmacokinetics, and an excellent safety profile^55–57^. Support for this strategy also comes from other treatments that likewise target this pathway, including intravenous infusions with hyaluronidase^58^ or anti-CD44 antibodies^23^. However, these approaches are impractical, immunosuppressive or toxic such that it has not been possible to target this pathway therapeutically^59, 60^.

## METHODS

### Donors and Procurement of Human Tissues

Pancreas tissue sections from 11 T2D and 17 non-diabetes control subjects were also obtained through the JDRF-sponsored Network for Pancreatic Organ Donors with Diabetes (nPOD) program. All tissues showed well-preserved morphology without any evidence of autolysis. This study was carried out with the approval of the Institutional Review Board (IRB) of the Benaroya Research Institute. Clinical characteristics of pancreatic tissue donors are presented in Table 1.

Human islets for research were provided by the Alberta Diabetes Institute Islet Core at the University of Alberta in Edmonton with the assistance of the Human Organ Procurement and Exchange (HOPE) program, Trillium Gift of Life Network (TGLN) and other Canadian organ procurement organizations. Islet isolation was approved by the Human Research Ethics Board at the University of Alberta (Pro00013094). All donors’ families gave informed consent for the use of pancreatic tissue in research. Purified islets were cultured in RPMI at 5.5 mM glucose supplemented with 10% FBS. Islet function (i.e., glucose-induced insulin secretion) was assessed by islet perfusion assay on the day of arrival as previously described^61^.

### Human islet Affinity Histochemistry and Immunohistochemistry

5 µm paraffin-embedded pancreas sections from T2DM and non-diabetes human subjects were stained for HA using a biotinylated HA-binding protein (b-HABP) prepared from cartilage and for immunohistochemistry as described previously^36^. Antibodies to synaptophysin (Dako, Carpinteria, CA) were used at dilutions 1:100. The secondary antibodies Alexa Fluor 488– or Alexa Fluor 555–conjugated IgG (Molecular Probes, Grand Island, NY) were used at 3 mg/mL. The nuclei were visualized with DAPI (Sigma-Aldrich, St. Louis, MO). Positive and negative controls were included in each staining experiment where tissues either expressed or lacked the marker of interest. Tissues were examined using a Leica DM IRB microscope, and images were acquired using a Spot Xplorer camera and imaging software. All the images were taken under the same experimental settings and light exposure. Images were analyzed using imageJ analysis software.

### Animals

Male B6.db/db LepR^-/-^ mice at (4 – 5 weeks) were purchased from Jackson Laboratories (B6.BKS(D)-Leprdb/J). The same mice were crossed with CD44^-/-^ mice, also bought from Jackson Laboratories (B6.129(Cg)-*Cd44^tm1Hbg^*/J) to generate db/dbxCD44^-/-^ mice. CD44fl/fl mice were crossed with INS1cre mice bought from Jackson Laboratories (B6(Cg)-Ins1^tm1.1(cre)Thor^/J) to generate CD44fl/flxINS1cre mice. Survivinfl/fl mice bought from Jackson Laboratories (B6.129P2^- Birc5tm1Mak^/J) were crossed with the INS1cre mice in order to generate the survivin/INS1cre mice. All mice were maintained in specific pathogen-free AAALAC-accredited animal facilities at Stanford University and handled in accordance with institutional guidelines.

### 4-MU and 4-MUG treatment

4-MU (Alfa Aesar, Ward Hill, MA) was pressed into the mouse chow by TestDiet^®^ as previously described^36^. We previously determined that this chow formulation delivers 250 mg/mouse/day, yielding a plasma drug concentration of 640.3 ± 17.2 nmol/L in mice, as measured by HPLC-MS. The 4-MUG (Chem-Impex International, Wood Dale, IL) was dosed in the drinking water at 2 mg/ml. Mice were initiated on the 4-MU chow or the 4-MUG drinking water at four to five weeks of age, unless otherwise noted, and were maintained on this diet until they were euthanized.

### Weight and diabetes monitoring

Beginning at four weeks of age, mice were weighed weekly as well as bled *via* the tail tip puncture for the determination of their blood glucose level (BG) using a blood glucose meter and blood glucose monitoring strips. When two consecutive blood glucose readings of 300 mg/dL were recorded, animals were considered diabetic.

### Flow cytometry and phenotyping

Mouse leukocyte populations were isolated from inguinal, mesenteric and pancreatic lymph nodes and spleens as previously described^59^. Mouse flow cytometry experiments used the following fluorochrome-labeled antibodies: CD44 (IM7) from BD- Biosciences (San Jose, CA). Flow cytometry protocols were done as previously described^60^. A FACSCaliber flow cytometer (BD, Franklin Lakes, NJ) was used to collect data. Analysis was performed using CELLQuest (BD) and FlowJo (Treestar Inc., Ashland, OR) software.

### Tissue processing and imaging

Tissues for histochemistry were taken out of the animals and immediately transferred into 10 % neutral buffered formalin (NBF) or methyl carnoys (MC) fixation. The tissue was processed to paraffin on a Leica ASP300 Tissue Processor (Leica Microsystems Inc., Buffalo Grove, IL). Then 5 µm thick sections were cut on a Leica RM 2255 Microtomes (Leica Microsystems Inc.).

For HA affinity histochemistry (AFC) the Bond Intense R Detection kit, a streptavidin-horse radish peroxidase (HRP) system, (Leica Microsystems, Inc.) was used with 4 µg/mL biotinylated-HABP in 0.1 % BSA-PBS as the primary. The Bond Polymer Detection Kit was used for all other immunohistochemistry. This detection kit contains a goat anti-rabbit conjugated to polymeric HRP and a rabbit anti-mouse post primary reagent for use with mouse primary antibodies.

The paraffin slides for the immune-fluorescence HA staining, were deparaffinized in xylene and were brought down to PBS in descending concentrations of ethanol. The slides were then rinsed several times in PBS, and blocked in 4 % BSA for 5 h. The tissues were probed with 4 µg/mL HABP in blocking medium, overnight at RT. The slides were rinsed in PBS for 30 min, before the secondary Streptavidin (S32354, Molecular Probes, Grand Island, NY) was used at 1:400 for 2 h. The slides were rinsed in PBS and the nuclei were stained with Propidium Iodide (P3566, Molecular Probes, Grand Island, NY) at 1:200. The Propidium Iodide was mixed into the mounting medium with Prolog Gold Anti-fade (P36930, Molecular Probes). CD44 IHC required pretreatment using heat-mediated antigen retrieval with EDTA at high pH (Bond Epitope Retrieval

Solution 2) for 10 minutes. For CD44 IHC, sections were incubated for 30 minutes with 0.5 µg/ml rat anti-CD44 clone IM7 (Thermo Scientific). For the survivin IHC a survivin rabbit mAB (Cell Signaling, Danvers, MA) was used at 1:200. For the TUNEL staining, a TUNEL Assay Kit (ab206386, Abcam, Cambridge, UK) was used according to the provided instructions.

All images were visualized using a Leica DMIRB inverted fluorescence microscope equipped with a Pursuit 4-megapixel cooled color/monochrome charge-coupled device camera (Diagnostic Instruments, Sterling Heights, MI). Images were acquired using the Spot Pursuit camera and Spot Advance Software (SPOT Imaging Solutions; Diagnostic Instruments). Image analysis was performed according using Image J (NIH), as described previously^61^.

### Blood glucose measurements, IPGTT, ITT

Mice were bled via the tail tip puncture for the determination of their blood glucose level using a blood glucose meter and blood glucose monitoring strips (Relion, Bayer). For IPGTT, mice were fasted 16 hours overnight and given i.p. 2 g of glucose/kg body weight in PBS. Blood glucose values were measured before and after glucose administration at 0, 15, 30, 60, and 120 minutes. The glucose meter used in this study has an upper threshold of 600 mg/dl. Therefore, values ≥600 mg/dl were diluted 2-fold and repeated.

### Serum Insulin ELISA

Blood was collected from mice via tail vein incision and using heparinized capillary tubes (BD Biosciences). The blood was centrifuged at 3000 g for 15 minutes at RT. The supernatant was collected and serum insulin levels were determined in triplicate using a rat/mouse insulin Enzyme-Linked Sorbent Assay (ELISA) kit from Millipore (EZRMI-13K) according to the instructions of the manufacturer. Rat insulin of 0–20 ng/ml was used as a standard.

### Mouse islet isolation and culture

To obtain mouse islets mice (age 12 – 14 weeks, n = 4/group) were euthanized and cervically dislocated immediately prior to dissection. Islets were prepared by injecting collagenase (10 ml of 0.23 mg/ml liberase; Roche Molecular Biochemicals, Indianapolis, IN) into the pancreatic duct and surgically removing the pancreas. The pancreases were placed into 15-ml conical tubes containing 5 ml of 0.23 mg/ml liberase and incubated at 37 degrees for 30 min. The digested islet mix was filtered (400 mm stainless steel screen), rinsed (Hank’s buffered salt solution), and purified in a gradient solution of Histopaque. Islets were cultured for 18–24 h in RPMI media 1640 supplemented with 10% heat inactivated FBS before further experimentation. All procedures were approved by the Stanford University Institutional Animal Care and Use Committee.

### Western blots

Min6 western blots were performed after the different treatments of 300 µM 4- MU and 100 µg/ml HA for 24 h and 48 h. Min6 cells were washed once with cold PBS and lysed with cell lysis buffer (50 mM Tris-HCl pH 8, 137 mM NaCl, 10% glycerol, 1% NP-40) supplemented with protease and phosphatase inhibitors (Millipore-Sigma). Protein concentrations was determined with a Bicinchoninic Acid Assay Kit (Pierce). Immunoblotting experiments were performed using 20 µg of total proteins per lane loaded on Precast SDS gels (Genscript). After electrophoresis, proteins were transferred to a hydrophilic PVDF membrane (Imobilon E, Millipore-Sigma). Membranes were blocked with TBS containing 1 % casein and 2 % BSA followed by incubation with primary antibodies (survivin 1:2000, 71G4B7, Cell Signaling; beta actin 1:5000 sc-4778, Santa Cruz Biotechnology; GLUT2 1:1000, 720238 Thermo Fisher Scientific) for 16 h at 4C in blocking buffer. After washing, membranes were incubated for 1 h with the respective secondary antibodies conjugated with NIR dies. Membranes were thoroughly washed and imaged on a Li-Cor Odyssey laser scanner (Li-Cor Biosciences). Signal quantification was performed using Image studio (Li-Cor Biosciences).

### Islet insulin response

Islet function was assessed by monitoring the insulin secretory response of purified islets after 300 µM 4-MU treatment. These were incubated in 3 mM glucose Krebs-Ringer bicarbonate (KBR) solution (2.6 mmol/l CaCl_2_/2H_2_O, 1.2 mmol/l MgSO_4_/7H_2_O, 1.2 mmol/l KH_2_PO_4_, 4.9 mmol/l KCl, 98.5 mmol/l NaCl, and 25.9 mmol/l NaHCO3 (all from Sigma-Aldrich, St. Louis, MO) supplemented with 20 mmol/l HEPES/NaHEPES (Roche Molecular Biochemicals, Indianapolis, IN) and 0.1 % BSA (Serological, Norcross, GA). 40 islets per condition were placed into a 96 well plate (10 islets per well in quadruplicate) containing 200 ml of 20 mM glucose KRB and incubated for 2 hours at 37^0^C and 5 % CO_2_. The supernatant was collected for insulin measurement. Insulin concentrations in these experiments were analyzed with an insulin ELISA kit as described above.

### RT-PCR

Min6.luc and Min6.CD44 cells were harvested for total RNA isolation using the High Pure RNA isolation kit (Roche Applied Science) and reverse-transcribed using the High Capacity cDNA Reverse Transcription kit (Applied Biosystems). Real-time quantitative polymerase chain reaction (qRT-PCR) quantification of insulin mRNA and 18S rRNA levels was performed using TaqMan Gene Expression Assays (Applied Biosystems). Data for mRNA expression are provided as mean ± standard error of the mean of the estimated copy number, normalized to 18S rRNA, and differences between diabetic and non-diabetic islets were analyzed.

### HA quantification

Samples were first lyophilized and weighed, then were digested with proteinase K (250 µg/mL) in 100 mM ammonium acetate pH 7.0 overnight at 60°C. After digestion, the enzyme was inactivated by heating to 100°C for 20 minutes. Total amount of HA was determined by a modified competitive ELISA in which the samples to be assayed were first mixed with biotinylated HA-binding protein (b-HABP) and then added to HA-coated microtiter plates, the final signal being inversely proportional to the level of hyaluronan added to the bPG.

### Measurement of cell supernatant levels of HA

Samples were thawed and then assayed for HA levels in a single batch using a modified HA Enzyme-Linked Sorbent Assay (ELISA). Each sample was analyzed in triplicate with a mean value obtained for each individual.

### Islet transplantation

Islets from healthy B6.CD44^+/+^ or B6.CD44^-/-^ were transplanted into autologous db/db recipients previously rendered diabetic using STZ. Three days prior to surgical implantation of BIs, db/db mice were treated with a high dose (200 mg/kg) of STZ made as a stock solution of 7.5 mg/ml STZ in 100 mM citrate, pH 4.2 (prepared and filtered at 0.22 μm immediately prior to intraperitoneal injection). Islet transplantation was performed under the kidney capsule using a previously published protocol^61^. Insulin was dosed with diluted Levimir daily until recipient mice were normo-glycemic (∼2 weeks).

### Statistical analysis

Human data are expressed as means ± SD of n independent measurements. Murine data are expressed as ± SE of n independent measurements, unless otherwise noted. The comparison between 2 groups was performed with unpaired *t* tests. Significance of the difference between the means of three groups of data was evaluated using the Mann-Whitney U test or one-way ANOVA, respectively. Correlation analysis was performed using the non-parametric Spearman correlation test. A p value less than <0.05 was considered statistically significant. Data analysis was performed with the use of GraphPad Prism 9.0 software.

## ACKNOWLEDGEMENTS

This work was supported in part by the National Institutes of Health grants R01 DK096087-01, R01 HL113294-01A1, and U01 AI101984 to PLB and R01 D088082 to RLH. This work was also supported by grants from the JDRF 3-PDF-2014-224-A-N to NN and 1-SRA-2018-518-S-B Innovation Award to PLB and by grants from the Harrington Institute, and Stanford SPARK to PLB, and by a grant from the Stanford Diabetes Research Center (SDRC) to NN. We thank the SDRC Stanford Islet Research Core (SIRC) and Diabetes Immune Monitoring Core (DIMC) which are part of the P30 award (P30DK116074) from the NIH/NIDDK.

## ABBREVIATIONS

4-MU: 4-methylumbelliferone
4-MUG: 4-methylumbelliferyl glucuronide
B6: C57Bl6 mice
b-HABP: biotinylated HA-binding protein
BG: blood glucose
ECM: extracellular matrix
eGWAS: expression-based genome-wide association study
HA: hyaluronan
HAS: HA synthases
HOPE: Human Organ Procurement and Exchange program
IPGTT: intra peritoneal glucose tolerance test
ITT: insulin tolerance test
JDRF: Juvenile Diabetes Research Foundation
nPOD: Network for Pancreatic Organ Donors
ST: staurosporine
STZ: streptozotocin
T2D: Type 2 diabetes
TGLN: Trillium Gift of Life Network
TMRE: tetramethylrhodamine ethyl ester

## SUPPLEMENTAL FIGURE LEGENDS

**Supplemental Figure 1. 4-MU has modest effects on BG and glucose responses in B6 mice. A.** Random fed blood glucose (BG) values over time for db/db mice treated with control chow and db/db mice treated with 4-MU chow. **B.** IPGTT performed on fasting db/db mice on control and 4- MU chow. n = 6-8 mice per group. SE is shown. None of these comparisons were statistically significant.

**Supplemental Figure 2. Mice initially eat less 4-MU chow.** Remaining chow in the cages was measured for a duration of 15 days and compared % chow remaining between control chow and 4-MU chow treated mice. Chow was refilled and set to 100% on day 1 and day 9. n = 20-25 mice per group.

**Supplemental Figure 3. 4-MUG prevents hyperglycemia in obese db/db mice. A-C.** Representative images (**A**), blood glucose (BG) values (**B**), and weights (**C**) over time for db/db mice treated with control chow, db/db mice treated with 4-MUG in drinking water as well as for db/+ heterozygous littermates treated with regular drinking water (control). * = p<0.05 db/db control vs db/+ control. ^ = p<0.05 db/db control vs db/db 4-MUG. n = >10 mice per group for each time point. **D, E.** Random (fed) BG values (**D**) and weights (**E**) for db/db mice treated with 4-MUG or placebo for 4 weeks as well as their db/+ littermate controls on control chow. Each dot in **D** and **E** represents one mouse. Values over 600 mg/dL were excluded. **F.** IPGTT performed on fasting db/db mice treated with and without 4-MUG. N = at least 8 mice per group for each figure panel. * = p<0.05.

**Supplemental Figure 4. The absence of CD44 contributes to modest effects on glycemic control in conventional B6 mice. A.** Random (fed) BG values for db/db.CD44^+/+^ and db/db.CD44^-/-^ mice at ages between 8 and 25 weeks of age. N = 50 mice per group. **B.** IPGTT performed on fasting db/db.CD44^+/+^ and db/db.CD44^-/-^ mice. N = 6-8 mice per group. * = p<0.05.

**Supplemental Figure 5. Both HA and CD44 adversely impact insulin mRNA levels in islets. A.** A murine β-cell line, Min6 cells (MIN6.luc) and Min6 engineered to overexpress CD44 (MIN6.CD44), were cultured in media containing glucose and their insulin mRNA expression was measured, normalized to 18S. B. MIN6 cells were cultured in media containing glucose, 100 µg/mL HA and 50 µg/mL 4-MU for 48 and 72 hours. Their insulin mRNA expression is normalized to 18S. Data are representative of 3 independent experiments. * = p<0.05.

**Supplemental Figure 6. The STZ receptor GLUT-2 expression is not altered by 4-MU or CD44. A.** Glut-2 expression of Min6 cells, control (Min6.Luc), Min6.CD44 overexpressing CD44 and Min6 control cells treated with 4-MU for 24 hours. **B**. Quantified GLUT-2 expression of Min6 cells in WB in **A**.

**Supplemental Figure 7. CD44 contributes to glycemic control in conventional B6 mice. A.** IPGTT performed on fasting db/db.CD44^+/+^ and db/db.CD44^-/-^ mice 21 days after STZ treatment. **B.** Random fed BG over time after mice were treated with high dose (200 mg/kg) STZ. n = 6-8 mice per group. * = p<0.05.

