## Supplemental Figures for "Inhibition of hyaluronan synthesis prevents β-cell loss in obesity-associated type 2 diabetes"

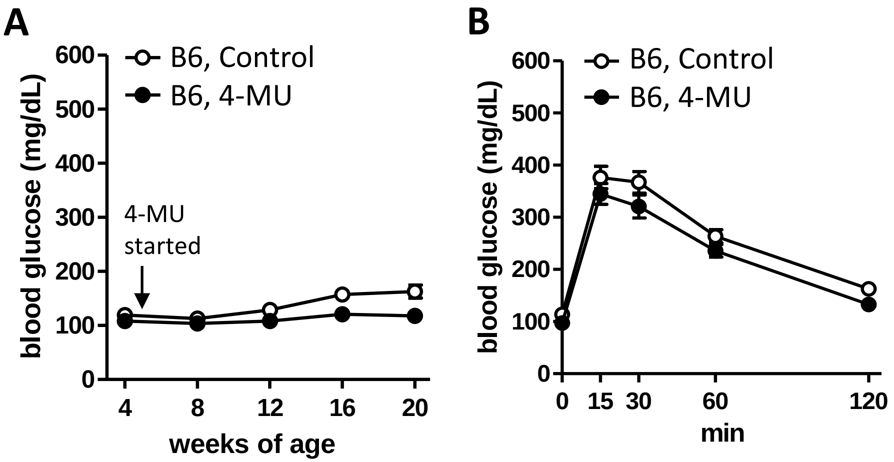


**Supplemental Figure 1. 4-MU has modest effects on BG and glucose responses in B6 mice. A.** Random fed blood glucose (BG) values over time for db/db mice treated with control chow and db/db mice treated with 4-MU chow. **B.** IPGTT performed on fasting db/db mice on control and 4-MU chow. n = 6-8 mice per group. SE is shown. None of these comparisons were statistically significant.

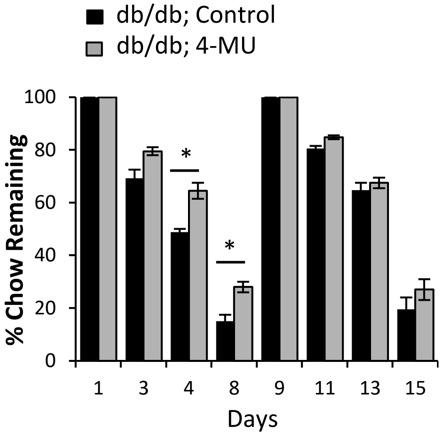


**Supplemental Figure 2. Mice initially eat less 4-MU chow.** Remaining chow in the cages was measured for a duration of 15 days and compared % chow remaining between control chow and 4-MU chow treated mice. Chow was refilled and set to 100% on day 1 and day 9. n = 20-25 mice per group.

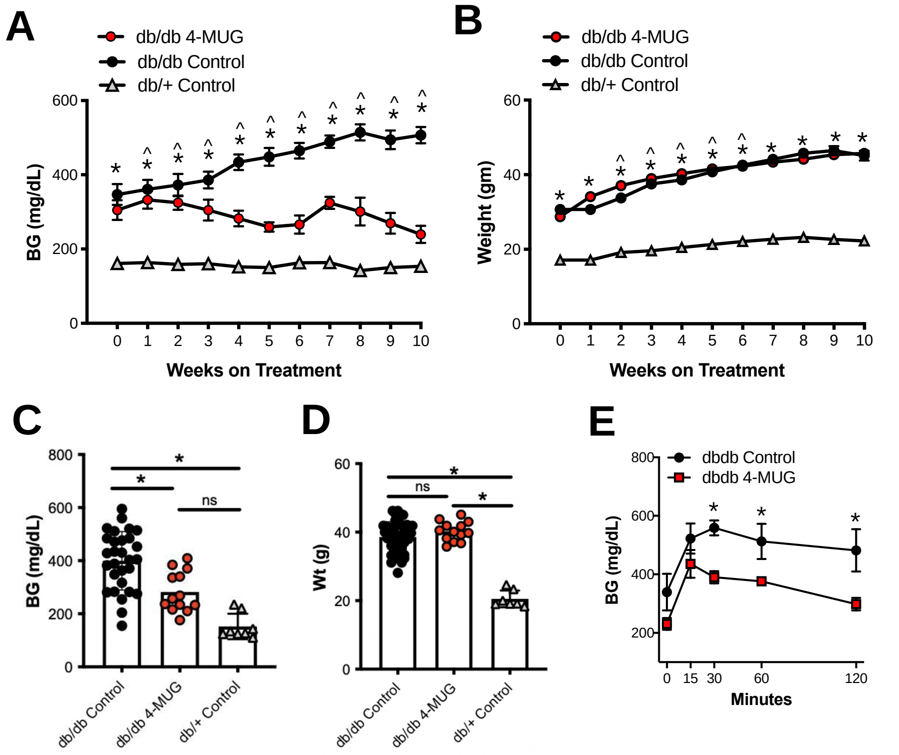


**Supplemental Figure 3. 4-MUG prevents hyperglycemia in obese db/db mice. A-C.** Representative images (**A**), blood glucose (BG) values (**B**), and weights (**C**) over time for db/db mice treated with control chow, db/db mice treated with 4-MUG in drinking water as well as for db/+ heterozygous littermates treated with regular drinking water (control). * = p<0.05 db/db control vs db/+ control. ^ = p<0.05 db/db control vs db/db 4-MUG. n = >10 mice per group for each time point. **D, E.** Random (fed) BG values (**D**) and weights (**E**) for db/db mice treated with 4-MUG or placebo for 4 weeks as well as their db/+ littermate controls on control chow. Each dot in **D** and **E** represents one mouse. Values over 600 mg/dL were excluded. **F.** IPGTT performed on fasting db/db mice treated with and without 4-MUG. N = at least 8 mice per group for each figure panel. * = p<0.05.

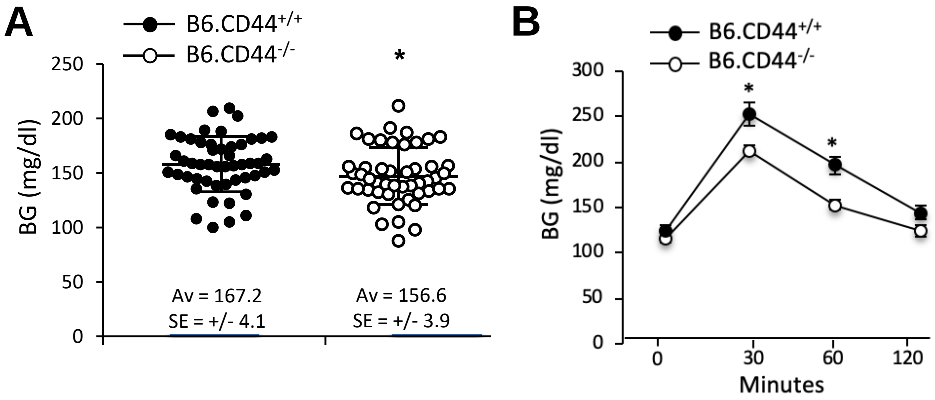


**Supplemental Figure 4. The absence of CD44 contributes to modest effects on glycemic control in conventional B6 mice. A.** Random (fed) BG values for db/db.CD44$^{+/+}$ and db/db.CD44$^{-/-}$ mice at ages between 8 and 25 weeks of age. N = 50 mice per group. **B.** IPGTT performed on fasting db/db.CD44$^{+/+}$ and db/db.CD44$^{-/-}$ mice. N = 6-8 mice per group. * = $p<0.05$.

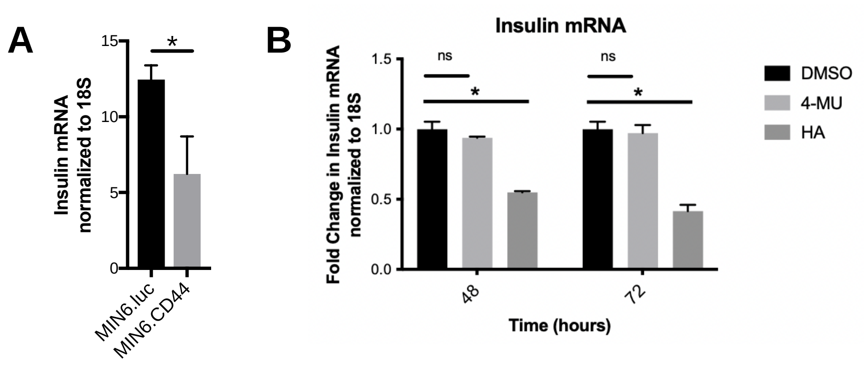


**Supplemental Figure 5. Both HA and CD44 adversely impact insulin mRNA levels in islets. A.** A murine β cell line, Min6 cells (MIN6.luc) and Min6 engineered to overexpress CD44 (MIN6.CD44), were cultured in media containing glucose and their insulin mRNA expression was measured, normalized to 18S. B. MIN6 cells were cultured in media containing glucose, 100 µg/mL HA and 50 µg/mL 4-MU for 48 and 72 hours. Their insulin mRNA expression is normalized to 18S. Data are representative of 3 independent experiments. * = $p<0.05$.

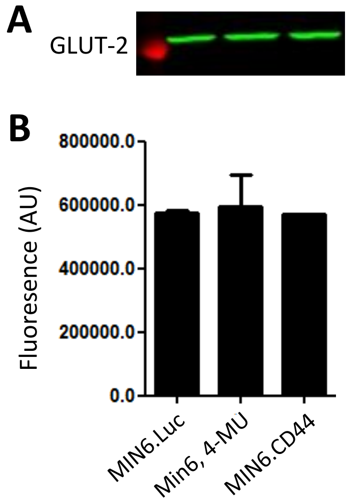


**Supplemental Figure 6. The STZ receptor GLUT-2 expression is not altered by 4-MU or CD44.** Glut-2 expression of Min6 cells, control (Min6.Luc), Min6.CD44 overexpressing CD44 and Min6 control cells treated with 4-MU for 24 hours. **B**. Quantified GLUT-2 expression of Min6 cells in WB in **A**.

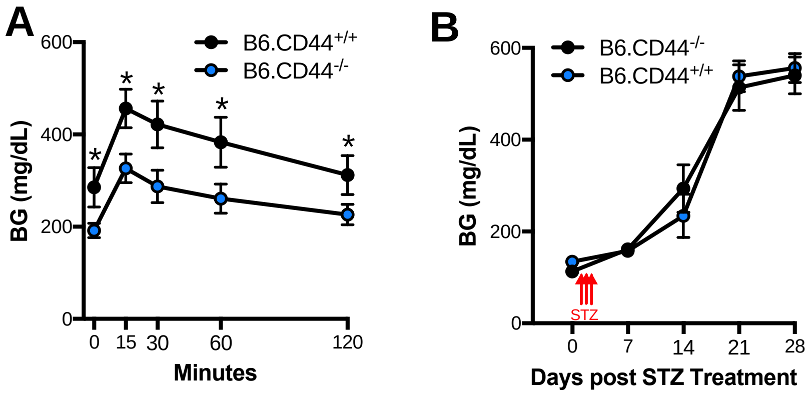


**Supplemental Figure 7. CD44 contributes to glycemic control in conventional B6 mice. A.** IPGTT performed on fasting db/db.CD44$^{+/+}$ and db/db.CD44$^{-/-}$ mice 21 days after low dose (40 mg/kg) STZ treatment. **B.** Random fed BG over time after mice were treated with high dose (200 mg/kg) STZ. n = 6-8 mice per group. * = $p<0.05$.
